# Genomes of novel Myxococcota reveal severely curtailed machineries for predation and cellular differentiation

**DOI:** 10.1101/2021.07.06.451402

**Authors:** Chelsea L. Murphy, R. Yang, T. Decker, C. Cavalliere, V. Andreev, N. Bircher, J. Cornell, R. Dohmen, C. J. Pratt, A. Grinnell, J. Higgs, C. Jett, E. Gillett, R. Khadka, S. Mares, C. Meili, J. Liu, H. Mukhtar, Mostafa S. Elshahed, Noha H. Youssef

**Author notes:** Corresponding author: Mailing address: Oklahoma State University, Department of Microbiology and Molecular Genetics, 1110 S Innovation Way, Stillwater, OK 74074.

## Abstract

Cultured Myxococcota are predominantly aerobic soil inhabitants, characterized by their highly coordinated predation and cellular differentiation capacities. Little is currently known regarding yet-uncultured Myxococcota from anaerobic, non-soil habitats. We analyzed genomes representing one novel order (o__JAFGXQ01) and one novel family (f__JAFGIB01) in the Myxococcota from an anoxic freshwater spring in Oklahoma, USA. Compared to their soil counterparts, anaerobic Myxococcota possess smaller genomes, and a smaller number of genes encoding biosynthetic gene clusters (BGCs), peptidases, one- and two-component signal transduction systems, and transcriptional regulators. Detailed analysis of thirteen distinct pathways/processes crucial to predation and cellular differentiation revealed severely curtailed machineries, with the notable absence of homologs for key transcription factors (e.g. FruA and MrpC), outer membrane exchange receptor (TraA), and the majority of sporulation-specific and A-motility-specific genes. Further, machine-learning approaches based on a set of 634 genes informative of social lifestyle predicted a non-social behavior for Zodletone Myxococcota. Metabolically, Zodletone Myxococcota genomes lacked aerobic respiratory capacities, but encoded genes suggestive of fermentation, dissimilatory nitrite reduction, and dissimilatory sulfate-reduction (in f_JAFGIB01) for energy acquisition. We propose that predation and cellular differentiation represent a niche adaptation strategy that evolved circa 500 Mya in response to the rise of soil as a distinct habitat on earth.

**Importance:** The Myxococcota is a phylogenetically coherent bacterial lineage that exhibits unique social traits. Cultured Myxococcoat are predominantly aerobic soil-dwelling microorganisms that are capable of predation and fruiting body formation. However, multiple yet-uncultured lineages within the Myxococcota has been encountered in a wide range of non-soil, predominantly anaerobic habitats; and the metabolic capabilities, physiological preferences, and capacity of social behavior of such lineages remains unclear. Here, we analyzed genomes recovered from a metagenomic analysis of an anoxic freshwater spring in Oklahoma, USA that represent novel, yet-uncultured, orders and families in the Myxococcota. The genomes appear to lack the characteristic hallmarks for social behavior encountered in Myxococcota genomes, and displayed a significantly smaller genome size and a smaller number of genes encoding biosynthetic gene clusters, peptidases, signal transduction systems, and transcriptional regulators. Such perceived lack of social capacity we confirmed through detailed comparative genomic analysis of thirteen pathways associated with Myxococcota social behavior, as well as the implementation of machine learning approaches to predict social behavior based on genome composition. Metabolically, these novel Myxococcota are predicted to be strict anaerobes, utilizing fermentation, nitrate rductio, and dissimilarity sulfate reduction for energy acquisition. Our result highlight the broad patterns of metabolic diversity within the yet-uncultured Myxococcota and suggest that the evolution of predation and fruiting body formation in the Myxococcoat has occurred in response to soil formation as a distinct habitat on earth.

## Introduction

The “Myxobacteria” represent a phylogenetically coherent lineage within the Domain Bacteria (1). Originally assigned to the class deltaproteobacteria (2, 3), they have recently been recognized as a separate phylum (Myxococcota) based on phylogenomic assessment; a proposal empirically supported by their distinct metabolic, and structural characteristics (4). The Myxococcota are highly social organisms, displaying specific behaviors (predation and fruiting body formation) that require a high level of kin recognition, cell-to-cell communication, and intercellular coordination (5). Indeed, cellular differentiation in the Myxococcota has aptly been described as the most successful foray for a prokaryotic organism into multicellularity (5).

Both predation and fruiting body formation processes involve differential gene activation and expression in seemingly equivalent cells, leading to distinct cellular differentiation and disparate fates in response to external environmental stimuli. As predators, model Myxococcota utilize an epibiotic strategy, where swarms of motile cells surround and lyse prey cells via the production of secondary metabolites and extracellular enzymes (6). Significant coordination of motility and lytic agents production between individual cells has been proposed as a means to enhance the efficiency of Myxococcota predator swarms ((7) but see (8)). Cellular differentiation entails the formation of elaborate multicellular structures (fruiting bodies) in response to nutrient depletion (9). The process involves cell aggregation and subsequent differentiation of a subset of cells into resistant myxospores, another subset into peripheral rods on the outside of the fruiting body adapted to rapidly respond to re-appearing nutrients, and a third subset undergoing programmed cell death (9, 10).

An extensive body of literature, spanning decades on the mechanistic basis of social behavior in model Myxococcota, is available (5, 6, 9, 11–13). Predation is enabled by two types of gliding motility (individual adventurous (A) motility and cooperative social (S) motility), exopolysaccharide production, secretion of secondary metabolites and extracellular enzymes (proteases and CAZymes), and mechanisms for kin/self-recognition and cheaters elimination (5) (this later is also important during aggregation and fruiting body formation). Starvation-induced cellular differentiation is mediated by four modules of highly interconnected and signal-responsive cascades of signal transduction networks that act sequentially as well as cooperatively to coordinate and time aggregation, fruiting body formation, and sporulation (11). Simultaneously, starvation-induced stringent response prompts the production of extracellular signals (most importantly A-signal, and C-signal) that activate a wide range of transcription factors to facilitate aggregation, regulate the onset of sporulation, and complete the development process (11).

Most cultured Myxococcota, henceforth referred to as “model Myxococcota”, are aerobic soil dwellers, known to inhabit the top layers of agricultural, forest, and even desert soils (14). This strong niche preference pattern attests to the contribution of their unique capacities to their success in soil ecosystems. The dual saprophytic/predatory capacities allow Myxococcota to utilize live microbial cells as well as microbial, floral, and faunal detritus as food sources (14). Their gliding motility allows them not only to get in close proximity with their prey, but also to access their insoluble substrates in soils (14). Their social behavior allows sharing resources (especially exoenzymes), enabling a more efficient process in which higher enzymatic activity is achieved when compared to individual cells (14). Fruiting bodies formation guarantee long term survival under adverse and highly fluctuating conditions in soil, as well as faster recovery and propagation under more favorable circumstances.

However, while Myxococcota appears to be most successful and prevalent in soils, members of this phylum have also been isolated from non-soil habitats (e.g. Nannocystaceae and Haliangiaceae from aerobic marine sediments (15, 16), and the facultative anaerobic *Anaeromyxobacter* from contaminated soils and sediments (17)). Further, multiple amplicon-based diversity surveys have identified Myxococcota-affiliated sequences in non-soil habitats, many of which represent anoxic/ hypoxic settings (18–24). Recently, the implementation of genome-resolved metagenomic approaches has resulted in the recovery of Myxococcota genomes from a wide range of non-soil habitats, almost invariably constituting a minor fraction of the population (25–30). Interestingly, many of these yet-uncultured lineages identified using 16S rRNA gene amplicon or metagenomic surveys appear to represent distinct novel, yet-uncultured lineages within the Myxococcota.

Given these observed strong niche adaptation patterns, as well as the predicted correlation between Myxococcota predation and cellular differentiation capacities, and successful propagation in soil, we hypothesized that novel, yet-uncultured Myxococota recovered from anaerobic non-soil habitats will display distinct metabolic, physiological, and lifestyle capacities when compared to their soil counterparts. Here, we analyzed multiple genomes representing two novel Myxococcota lineages recovered from the completely anoxic, sulfide-laden (8-10 mM) source sediment in Zodletone spring, an anaerobic sulfide-and sulfur-rich spring in Oklahoma. We investigated the metabolic capacities, physiological preferences, structural features, and potential social behavior of these lineages, and compared their predicted capacities to model social aerobic Myxococcota. Our results suggest that non-soil Myxococcota possess severely curtailed pathways for the typical social behavior of soil Myxococcota, potentially utilize fermentation and/or sulfate-reduction for energy generation as opposed to aerobic respiration, and show preferences to polysaccharide and sugars, rather than proteins and amino acids as carbon and energy sources. We argue that such differences provide important clues to the evolution of social behavior in the Myxococcota in light of our understanding of the history of soil formation and oxygen accumulation in the atmosphere.

## Materials and Methods

### Site description and geochemistry

Zodletone spring is located in the Anadarko Basin of western Oklahoma (N34.99562° W98.68895°). The site geochemistry has been previously described in detail (31–33). Briefly, at the spring source, sulfide and gaseous hydrocarbon-saturated waters are slowly (8 L/min) ejected, along with sediments that deposit at the source of the spring. High (8-10 mM) sulfide concentrations maintain complete anoxic conditions (oxygen levels < 0.1 μM) in the spring sediments. Oxygen concentrations in the 50 cm water column overlaying the sediments vary from 2–4 μM at the 2 mm above the source to complete oxygen exposure on the top of the water column (31).

### Sampling and nucleic acid extraction

The sampling and DNA extraction process have been previously described in detail (34, 35). Briefly, ten different sediment samples (≈50 grams each) were collected at 5-cm depth, as well as from the standing overlaid water in sterile containers. DNA was extracted from 0.5 grams of source sediments from each replicate sample. For water samples, 10L of water was filtered on 0.2 µm sterile filters, and DNA was directly extracted from filters. Extraction was conducted using the DNeasy PowerSoil kit (Qiagen, Valencia, CA, USA) according to manufacturer protocols.

### Metagenome sequencing, assembly, and binning

All extractions from sediment or water samples were pooled, and the pooled DNA was used for the preparation of sequencing libraries using the Nextera XT DNA library prep kit (Illumina, San Diego, CA, USA) as per manufacturer’s instructions. DNA sequencing was conducted using two lanes on the Illumina HiSeq 2500 platform and 150-bp pair-end technology for each of the water and sediment samples using the services of a commercial provider (Novogene, Beijing, China). Metagenomic sequencing of the sediments and water samples yielded 281 Gbp and 323 Gbp of raw data, respectively. Reads were assessed for quality using FastQC followed by quality filtering and trimming using Trimmomatic v0.38 (36). High quality reads were assembled into contigs using MegaHit (v.1.1.3) (37) with minimum Kmer of 27, maximum kmer of 127, Kmer step of 10, and minimum contig length of 1000 bp. Bowtie2 was used to calculate the percentage of reads that assembled into contigs and sequencing coverage for each contig. Contigs >1 Kbp were binned into draft metagenome-assembled genomes (MAGs) using Metabat (38) and MaxBin2 (39), followed by selection of the highest quality bins using DasTool (40). CheckM (41) was used for estimation of genome completeness, strain heterogeneity, and contamination by employing the lineage-specific workflow (lineage_wf flag). Quality designation of draft genomes was based on the criteria set forth by MIMAGs (42).

### Genomes classification

Taxonomic classifications followed the Genome Taxonomy Database (GTDB) release r95 (43), and were carried out using the classify_workflow in GTDB-Tk (v1.3.0) (44). Phylogenomic analysis utilized the concatenated alignment of a set of 120 single-copy bacterial genes (43) generated by the GTDB-Tk. Maximum-likelihood phylogenomic tree was constructed in RaxML using the PROTGAMMABLOSUM62 model and default parameters (45) and members of the Bdellovibrionota as an outgroup. To further assign genomes to putative families and genera, average amino acid identity (AAI), and shared gene content (SGC) were calculated using the AAI calculator [http://enve-omics.ce.gatech.edu/]. The arbitrary AAI cutoffs used were 49%, 52%, 56%, and 68% for class, order, family, and genus, respectively (46, 47). Further, Relative Evolutionary Divergence (RED) values, based on placement in the GTDB backbone tree (available at https://data.gtdb.ecogenomic.org/releases/release95/95.0/), were used to confirm the novelty of lineages to which the genomes are assigned. Values between 0.62 and 0.46 are indicative of a novel order, and values between 0.62 and 0.77 of a novel family.

### Annotation and genomic analysis

Protein-coding genes were predicted using Prodigal (48). GhostKOALA (49) was used for the annotation of every predicted open reading frame in bins and to assign protein-coding genes to KEGG orthologies (KOs), followed by metabolic pathways visualization in KEGG mapper (50). In addition, all genomes were queried with custom-built HMM profiles for alternate complex III components and hydrogenases. To construct HMM profiles, a representative protein was queried against the KEGG genes database using Blastp, and hits with e-values <1e^-80^ were downloaded, aligned, and used to construct an HMM profiles using the hmmbuild function of HMMer (v 3.1b2) (51). Hydrogenases HMM profiles were built using alignments downloaded from the Hydrogenase Database (HydDB) (52). The hmmscan function of HMMer (51) was used with the constructed profiles and a thresholding option of -T 100 to scan the protein-coding genes for possible hits. Further confirmation was achieved through phylogenetic assessment and tree building procedures. The 5S, 16S, and 23S rRNA sequences were identified using Barrnap 0.9 (https://github.com/tseemann/barrnap). tRNA sequences were identified using tRNAscan-SE (v 2.0.6, May 2020) [82]. Genomes were mined for CRISPR and Cas proteins using the CRISPR/CasFinder [83]. Proteases, peptidases, and protease inhibitors were identified using Blastp against the MEROPS database (53), while carbohydrate active enzymes (CAZymes) were identified by searching all ORFs from all genomes against the dbCAN hidden Markov models V9 (54) (downloaded from the dbCAN web server in September 2020). AntiSMASH 3.0 (55) was used with default parameters to predict biosynthetic gene clusters in the genomes. Metabolic reconstruction of reference Myxococcota type species genomes was obtained from the KEGG genomes database (https://www.genome.jp/kegg/genome/) and used for comparative genomics to Zodletone Myxococcota genomes.

### Phylogenetic analysis of dissimilatory sulfite reductase DsrAB

Predicted dissimilatory sulfite reductase subunits A and B were compared to reference sequences for phylogenetic placement by first aligning to corresponding subunits from sulfate-reducing taxa using Mafft (56). DsrA and DsrB alignments were concatenated in Mega X (57), and used to construct maximum-likelihood phylogenetic trees using FastTree (v 2.1.10) (58).

### Machine learning approaches

Genomes of type species of cultured social (i.e. experimentally verified to involve in predation and observed to undergo cellular differentiation) Myxococcota lineages (n=24) and all cultured non-social Myxococcota lineages (n=13) were downloaded from GenBank (June 2021). Lineages were assigned their “social” status using prior culture-based observations (15–17, 59). Genomes were annotated with KO numbers using GhostKOALA (49) using default parameters, and gene counts were assembled into a matrix. Informative KOs (n=634) were selected using indicator analysis, with the R package indicspecies (60) using the multipattern function, and used to build a predictive model. Data was then centered, scaled, and Box-Cox transformed using the R package caret {https://topepo.github.io/caret/}. Random forest classification training was performed in Python3 with the ensemble method of scikit-learn (v 0.24.1) (61). The data was randomly divided, with 75% of the data selected to serve as the training set and the remaining 25% reserved for model verification. Model training was performed with default parameters and 1000 estimators. The model successfully predicted the social behavior (with 100% accuracy) in the 25% data subset reserved for model verification. Social abilities of novel Myxococcota lineages were then predicted using the constructed model. Matthew’s Correlation Coefficient (62) was used to quantify classification accuracy. Including only pure cultures genomes’ in our model with experimentally verified social behavior ensured that the model was accurately trained on possession of social behavior rather than arbitrary genomic artifacts due to phylogenetic relatedness.

### Sequence and MAG accessions

The individual assembled Myxococcota MAGs analyzed in this study have been deposited at DDBJ/ENA/GenBank under the accessions JAFGVO000000000, JAFGQN000000000, JAFGWT000000000, JAFGTB000000000, JAFGXQ000000000, and JAFGIB000000000.

## Results

### Novel Myxococcota in Zodletone spring sediments

Six genomes were recovered from the anoxic black sediment sources (3 MAGs) and the water column (3 MAGs) of Zodletone spring, with estimated completion and contamination percentages ranging between 89.08-96.01% and 1.94-3.87%, respectively. Phylogenomic analysis (Fig 1), as well as AAI and RED values (Table 1) placed these genomes into one novel order (order JAFGXQ01; n=5), and one novel family (family JAFGIB01, order Polyangiales; n=1) within the class Polyangia. Intra-order AAI values assigned the five genomes in novel order JAFGXQ01 into two families (novel families JAFGVO01, JAFGXQ01), and three genera (novel genera JAFGVO01, JAFGQN01, JAFGXQ01). Names were assigned based on the assembly accession number of the most complete genome within each lineage (Tables 1 and S1).

**Figure 1.**
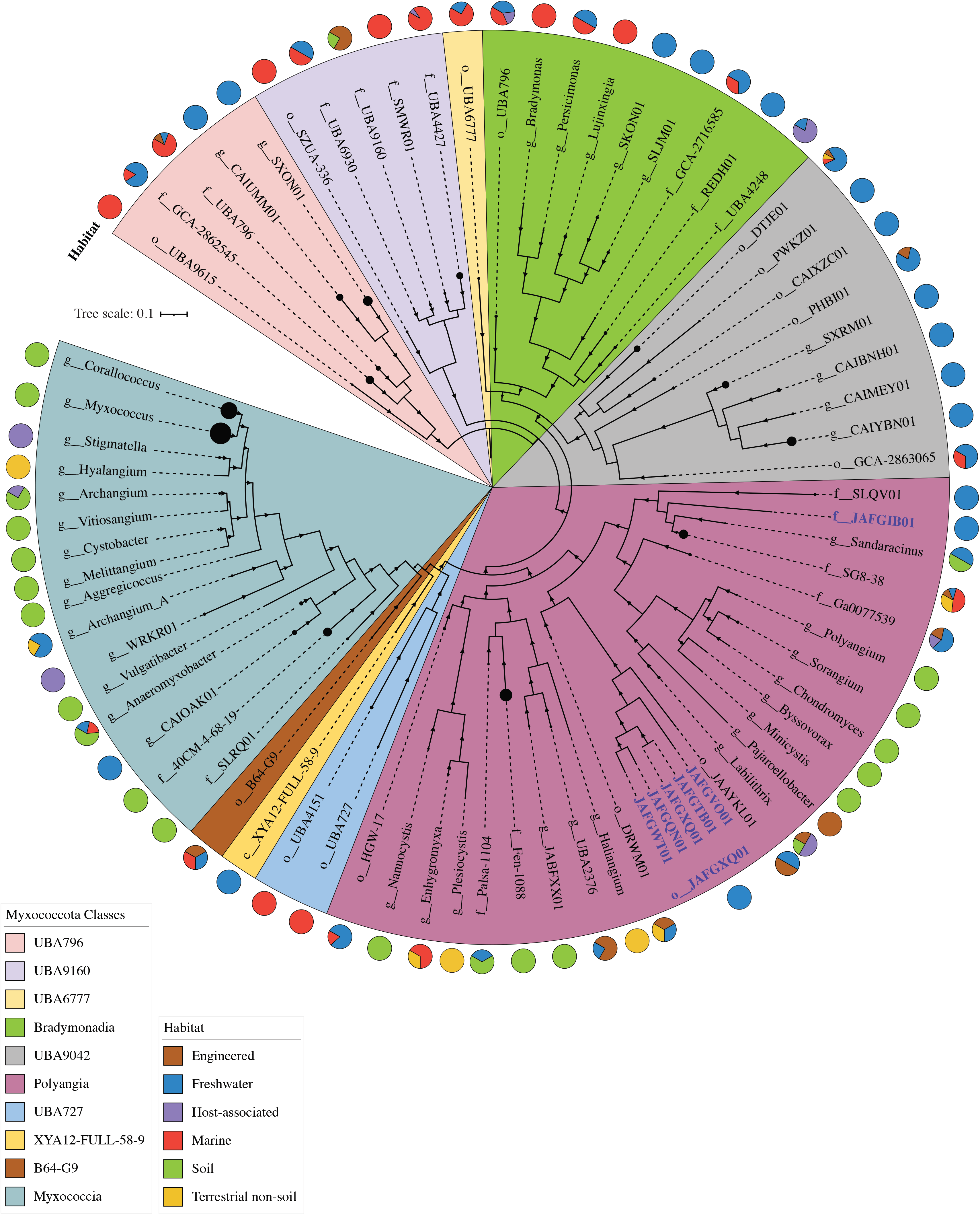
Phylogenomics of the Myxococcota including novel lineages from Zodletone spring. The maximum likelihood trees were constructed in RAxML (45) using all Myxococcota genomes available from GTDB r95 database based on the concatenated alignments of 120 housekeeping genes obtained from GTDB-Tk (44). The tree was rooted (root not shown) with the two Bdellovibrionota genomes *Halobacteriovorax marinus* (GCF_000210915.2) and *Bdellovibrio bacteriovorax* (GCF_000196175.1). The tree is wedged (shown as black circles at the end of branches) to represent genus level taxonomy (g__), unless the number of available genomes per genus is less than 5, in which case the family level (f__), or order level (o__) taxonomy is shown instead. The size of the wedge is proportional to the number of genomes. Bootstrap support values based on 100 replicates are shown as triangles for nodes with >70% support. Class-level taxonomy is color coded as shown in the legend. The track around the tree represents the ecosystem classification of the habitat from which the genomes originated. Zodletone genomes are labeled in blue bold text with their GenBank assembly accession number.

**Table 1.** Similarity statistics and GTDB classification of the MAGs analyzed in this study.

| Bin name | Assembly accession number | % completeness | % contamination | GTDB classification |  |  | Genus | Similarity statistics |  |  |
| --- | --- | --- | --- | --- | --- | --- | --- | --- | --- | --- |
|  |  |  |  | Class | Order | Family |  | RED<sup>1</sup> value | AAI<sup>2</sup> | SGC<sup>3</sup> |
| ZodW_Metabat.174 | JAFGVO01 | 91.24 | 3.26 | Polyangia | JAFGXQ01 | JAFGVO01 | JAFGVO01 | 0.490 | 38.1 ± 0.9 | 40.1 ± 5.62 |
| Zod_Metabat.76 | JAFGQN01 | 94.84 | 3.87 |  |  | JAFGXQ01 | JAFGQN01 | 0.493 |  |  |
| ZodW_Metabat.284 | JAFGWT01 | 89.08 | 3.87 |  |  |  |  | 0.489 |  |  |
| Zod_Metabat.947 | JAFGTB01 | 94.19 | 3.23 |  |  |  | JAFGXQ01 | 0.491 |  |  |
| ZodW_Metabat.4 | JAFGXQ01 | **96.01** | **3.23** |  |  |  |  | 0.491 |  |  |
| Zod_Metabat.169 | JAFGIB01 | **90.7** | **1.94** |  | Polyangiales | JAFGIB01 | JAFGIB01 | 0.660 | 40.1 ± 1.7 | 40.7 ± 5.07 |

<sup>1</sup>RED value: Relative Evolutionary Divergence values, based on placing the genomes in the Gtdb backbone bacterial tree (available at https://data.gtdb.ecogenomic.org/releases/release95/95.0/). RED values were used to confirm the novelty of the taxonomic lineage to which the genomes are assigned. Genomes are assigned to novel orders if their RED values fall between 0.62 and 0.46, and novel families if their RED values fall between 0.62 and 0.77.
<sup>2</sup>AAI: average amino acid identity calculated using the AAI calculator [http://enve-omics.ce.gatech.edu/]). The arbitrary AAI cutoffs used were 49%, 52%, 56%, and 68% for class, order, family, and genus, respectively.
<sup>3</sup>SGC: shared gene content

### Comparative genomic analysis between Zodletone Myxococcota and model Myxococcota

Comparative genomic analysis between Zodletone Myxococcota MAGs and genomes of all described type species in the phylum Myxococcota (n=27) was conducted. These genomes belong to classes Myxococcia (n=12), Polyangia (n=12), and Bradymonadia (n=3), and 20 of which exhibit the distinct social behavior of the Myxococcota (5-9, 11-13, 15-17, 59). We utilized only genomes from type species to ensure the availability of experimental data regarding various aspects of their lifestyle. Compared to model Myxococcota genomes (i.e. those shown to exhibit predation behavior and to form fruiting bodies, n=20); Zodletone genomes were significantly smaller (6.15 ± 1.28 Mb versus 11.44 ±2.52 Mb), with a lower gene count (5129±1005 versus 9461±1611) and GC content (49.53± 5.67 versus 69.68± 1.74) (Student t-test p-value <0.00001) (Figure 2). Further, multiple additional differences were observed in gene families previously implicated in mediating Myxococcota social lifestyle between Zodletone MAGs and genomes of social Myxococcota. Extracellular proteolytic enzymes are crucial components of the predatory machinery in Myxococcota, aiding in degrading prey-released proteins and/or inducing prey lysis (6). Zodletone genomes encoded a significantly lower number of proteases/peptidases when compared to model Myxococcota (58 ± 3.4 versus 130 ±17) (Figure 2, Table S2). Of note is the absence of representatives of MEROPS Family M15 (peptidoglycan endopeptidases) specifically implicated in prey cell lysis (Table S2). Further, model Myxococcota also secrete a plethora of secondary metabolites such as pigments, siderophores, bacteriocins, and antibiotics that attack and lyse their prey (6). Zodletone Myxococcota genomes encoded a significantly lower number of biosynthetic gene clusters (BGCs, 8±3), mostly belonging to the NRPS_PKS type. By comparison, model Myxococcota encoded a larger number of BGCs (38 ± 16), belonging to a wider range (NRPS, PKS, terpenes, siderophores, and phenazines) of BGC classes. (Figure 2, Table S3).

**Figure 2.**
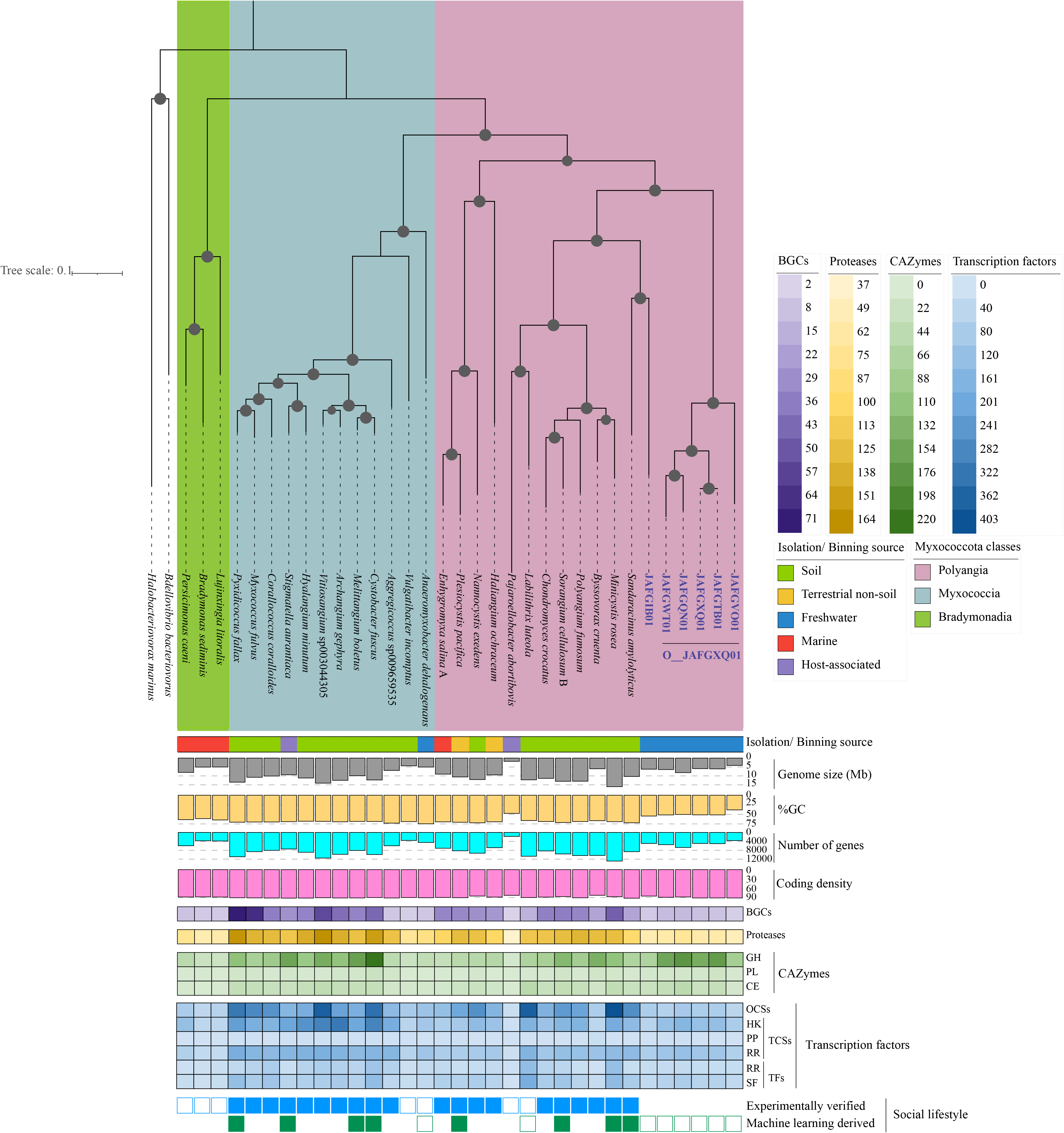
Comparative genomics of Zodletone novel Myxococcota genomes to the genomes of 27 type species belonging to the classes Myxococcia, Polyangia, and Bradymonadia. GenBank assembly accession numbers are: *Anaeromyxobacter dehalogenans* 2CP-1, GCF_000022145.1; *Haliangium ochraceum* DSM 14365, GCF_000024805.1; *Plesiocystis pacifica* SIR-1, GCF_000170895.1; *Corallococcus coralloides* DSM 2259, GCF_000255295.1; *Cystobacter fuscus* DSM 2262, GCF_000335475.2; *Hyalangium minutum*, GCF_000737315.1; *Sandaracinus amylolyticus*, GCF_000737325.1; *Archangium gephyra*, GCF_001027285.1; *Chondromyces crocatus*, GCF_001189295.1; *Vulgatibacter incomptus*, GCF_001263175.1; *Labilithrix luteola*, GCF_001263205.1; *Minicystis rosea*, GCF_001931535.1; *Melittangium boletus* DSM 14713, GCF_002305855.1; *Nannocystis exedens*, GCF_002343915.1; *Bradymonas sediminis*, GCF_003258315.1; *Lujinxingia litoralis*, GCF_003260125.1; *Polyangium fumosum*, GCF_005144585.1; *Persicimonas caeni*, GCF_006517175.1; *Myxococcus fulvus*, GCF_007991095.1; *Pyxidicoccus fallax*, GCF_012933655.1; *Stigmatella aurantiaca*, GCF_900109545.1; *Vitiosangium*, GCF_003044305.1; *Aggregicoccus*, GCA_009659535.1; *Pajaroellobacter abortibovis*, GCF_001931505.1; *Byssovorax cruenta*, GCA_001312805.1; *Enhygromyxa salina*, GCF_002994615.1; *Sorangium cellulosum* B, GCF_000067165.1. Zodletone genomes are labeled in blue bold text with their GenBank assembly accession number. Class-level taxonomy is color coded as shown in the legend. The tracks underneath the tree show: the ecosystem classification of the habitat from which the genomes originated, the assembly genome size (grey bars), GC content (yellow bars), total number of genes in the genome (cyan bars), and coding density (pink bars). The number of biosynthetic gene clusters (BGCs in purple), proteases (in yellow), CAZymes (in green), and transcription factors (in blue) encoded in each genome are shown as a heatmap with the color tone explained in the respective legend. Following the heatmaps, the two outermost tracks denote the presence (filled squares)/ absence (empty squares) of the Myxococcota typical social lifestyle as evidenced by experimental pure culture work (blue), and/or the machine learning approach (green) we used for lifestyle prediction based on the informative set of KO numbers provided in Dataset 1. Abbreviations: BGCs, biosynthetic gene clusters; GH, glycosyl hydrolases; PL, polysaccharide lyases; CE, carbohydrate esterase; OCSs, one-component systems; TFs, transcription factors; RR, response regulator; SF, sigma factor; TCSs, two-component systems; HK, histidine kinases; and PP, phospho-relay proteins.

Predation in model soil Myxococcota is also associated with secretion of extracellular or outer membrane CAZymes for targeting prey cell walls. While the overall numbers of CAZymes encoded in Zodletone genomes were not significantly different from those encountered in model soil Myxococcota genomes (Figure 2), the CAZyome families were significantly different between the two groups (Table S4). Specifically, model Myxococcota genomes were enriched in two GH families: GH23 (peptidoglycan lyases, consistent with their ability to target prey cell walls), and GH13 amylases (Student t-test p-value <0.02). Instead, Zodletone genomes were significantly (Student t-test p-value <0.02) enriched in GH and PL families targeting polysaccharide degradation, e.g. GH12, GH5, GH45, GH8, GH9 endoglucanases and cellobiohydrolases for cellulose degradation; GH10, GH11 xylanases for hemicellulose backbone degradation, and GH43 and GH54 xylosidases for hemicellulose side chain sugar removal; and PL1 and PL11 pectin/pectate/rhamnogalacturonan lyases for pectin degradation.

Finally, the collective social behavior in model soil Myxococcota is underpinned by an expanded arsenal of transcriptional factors. These include signal transduction one-component systems (OCS) (with a sensory domain and a response effector domain present in the same gene) and two-component systems (TCS) (with a sensor histidine kinase (HK), a partner response regulator (RR), and occasionally a phosphotransfer protein (PP)), as well as other transcriptional factors (TFs) including transcriptional regulators (TRs), and alternative sigma factors (SFs)”(63). Model soil Myxococcota genomes encode 241 ± 87 OCS genes, 329 ± 95 TCS genes, and 127 ± 56 TF genes. In contrast, Zodletone Myxococcota genomes encoded significantly lower OCSs, TCSs, and TFs (65±14, 198±58, and 69±18, respectively) (Student t-test p<0.05) (Figure 2, Table S5). This pattern of curtailed transcription factor repertoire in Zodletone genomes was pronounced in OCS and TCS systems (Figure 2), specifically OCS families AraC, ArsR, GntR, LysR, MarR, TetR, and Xre, and TCS-RR belonging to the families CheY, NarL, OmpR, and FrzZ (Table S5) (Student t-test p-value <0.05).

### Comparative genomics analysis of predation and cellular differentiation genes/pathways in the Myxococcota

We assessed the distribution patterns of pathways implicated in Myxococcota social behavior in Zodletone Myxococcota and compared them to *Myxococcus xanthus*, the model myxobacterium extensively studied for its social behavior, as well as to *Anaeromyxobacter dehalogenes*, *Vulgatibacter incomptus*, and *Labilithrix luteola*. These three isolates share many genomic features with social Myxococcota (large genome size, high %GC, large number of genes), but lack predation and fruiting body formation capacities (64, 65). Thirteen pathways were examined: four gene regulatory networks governing sporulation, aggregation, and fruiting body formation; exopolysaccharide production genes; two extracellular signals production gene clusters (A-signal and C-signal); aggregation, sporulation, and fruiting body formation genes; chemosensory pathways; developmental timers; two motility gene clusters (A-motility and S-motility); and outer membrane exchange genes (Figure 3, Tables 2, S. text and S6). Detailed analysis of the distribution patterns of genes in these pathways, as well as a background on their known functions are presented in the supplementary document (S. text). Collectively, our analysis clearly demonstrates that social behavior pathways were severely curtailed in Zodletone Myxococcota genomes (Tables 2, S. text, S6), where homologues of genes specific for the model Myxococcota social lifestyle (e.g. sporulation, extracellular signal production, motility, outer membrane exchange) were missing from Zodletone genomes. Specifically, the most notable deficiencies are the absence of homologues for extracellular signals production that control early events in aggregation, the absence of homologues for the two transcription factors FruA and MrpC that work cooperatively to control the start of sporulation (66–68), the absence of homologues for sporulation specific genes (11, 66, 68–70), and A motility-specific genes, and the absence of homologues for the outer membrane exchange receptor TraA that recognizes kin and allow membrane fusion (12) in Zodletone Myxococcota genomes or genomes of *Anaeromyxobacter dehalogenes. Labilithrix luteola,* and *Vulgatibacter incomptus* known from pure culture work not to aggregate into mounds, form fruiting bodies, or sporulate. Further, for the pathways with homologues identified, the majority of genes encoded proteins with domain similarities to known transcriptional response regulators or serine/ threonine kinases, peptidase domains, guanylate cyclase domains, chemotaxis-associated domains, or type IV pili, all of which are widely distributed in bacteria.

**Figure 3.**
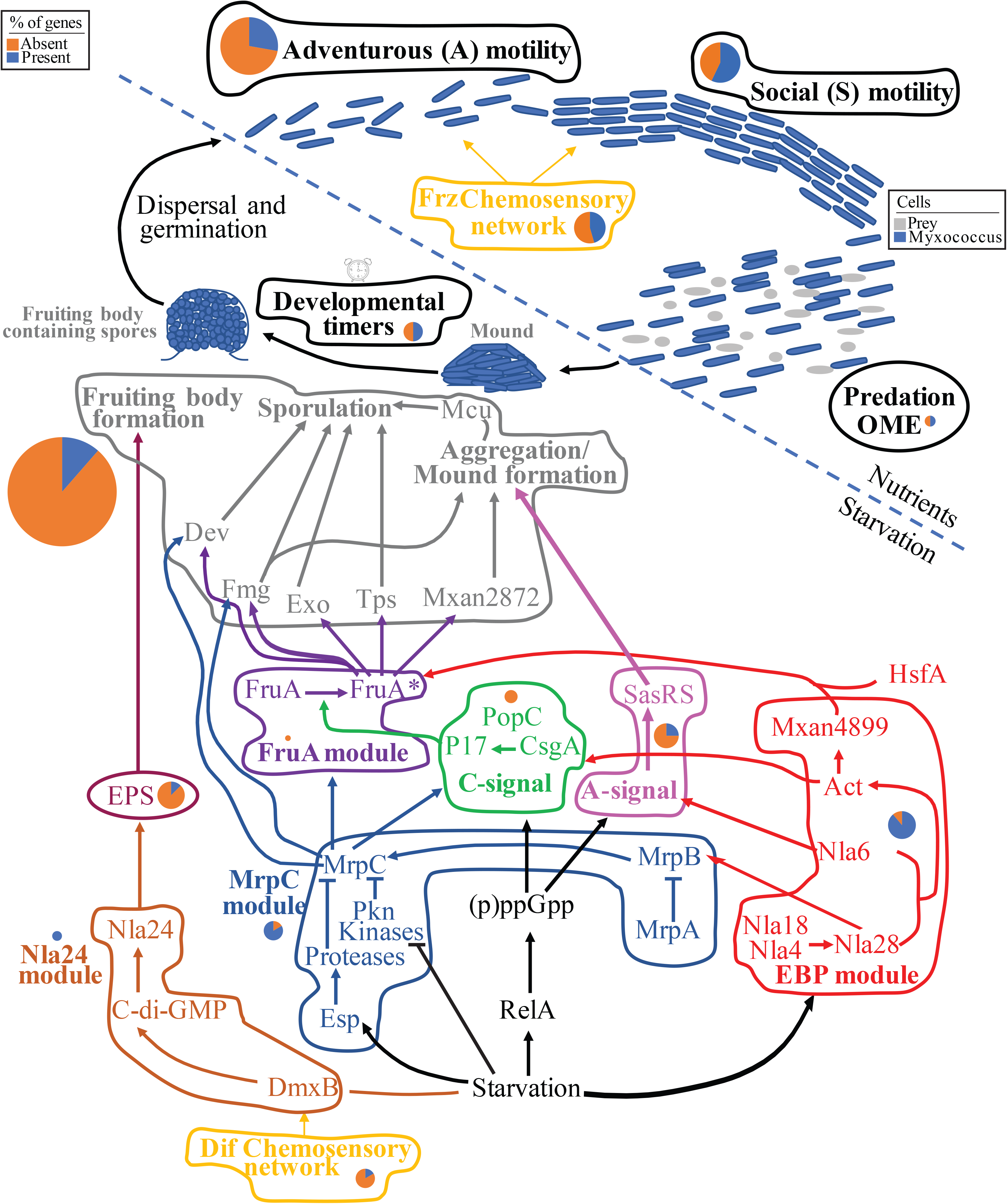
A cartoon depicting the thirteen pathways associated with Myxococcota social lifestyle examined in detail in this study. *Myxococcus* cells are shown as blue rods, while prey cells are depicted as grey cocci and rods. Pathways active during nutrient availability are shown above the dotted lines, while those induced by starvation are shown below the dotted line. Each of the pathways is shown in bold text within a color-coded outline. The same color code is used for the group of genes in each pathway and in Table S6. For each pathway, a pie-chart for the number of gene homologues identified in Zodletone genomes as a percentage of the total number of genes in the pathway is shown in blue, while the % of gene homologues absent are shown in orange. The size of the pie-chart is proportional to the number of genes in each pathway, and ranges from 1 (FruA module) to 35 (the aggregation/ sporulation/ fruiting body formation module). Arrow heads depict the effect where activation is shown as triangular arrowheads, and inhibition is shown as horizontal line arrowheads. FruA* denotes the active form of FruA. Abbreviations: EPS, exopolysaccharide; OME, outer membrane exchange.

**Table 2.**
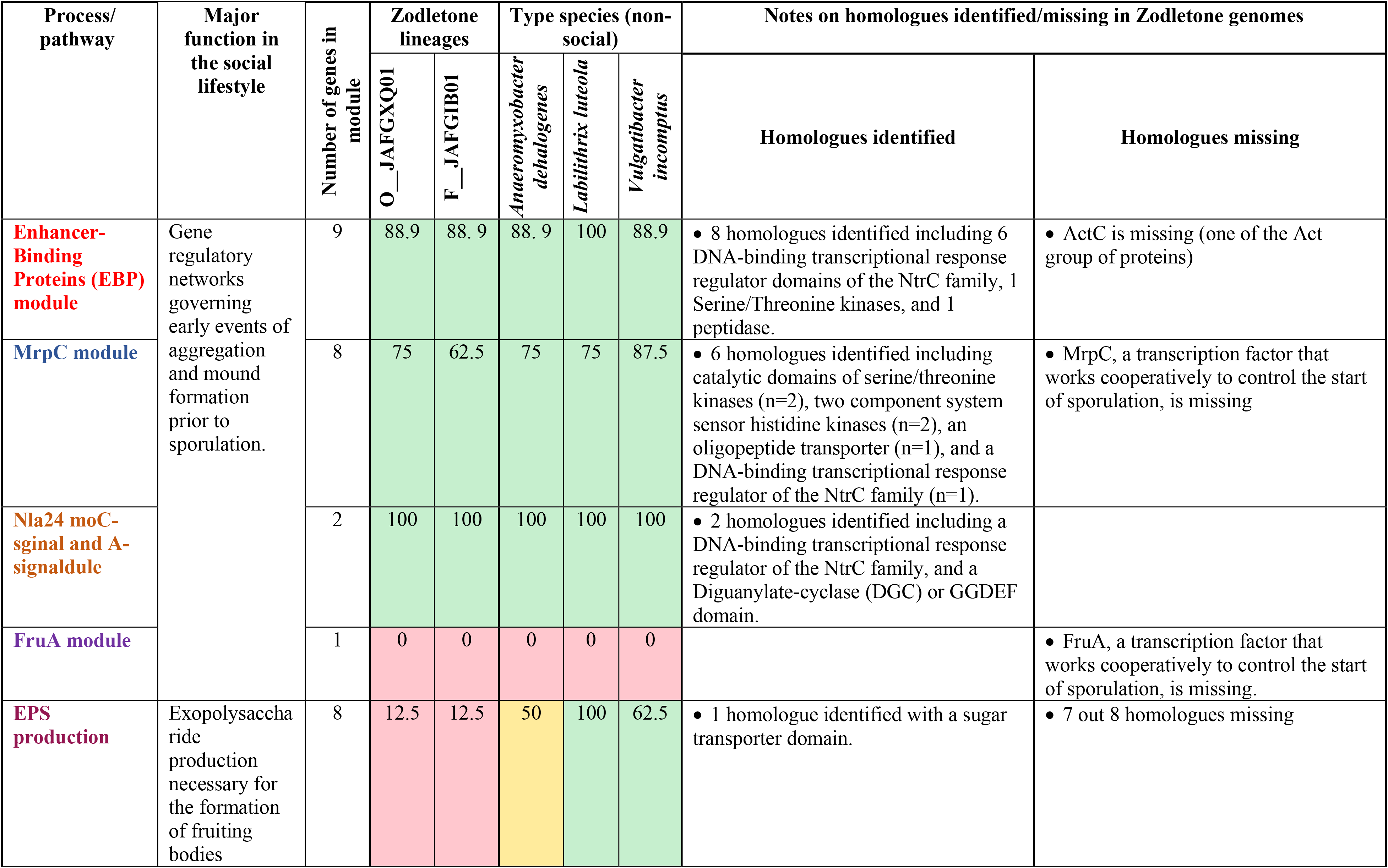
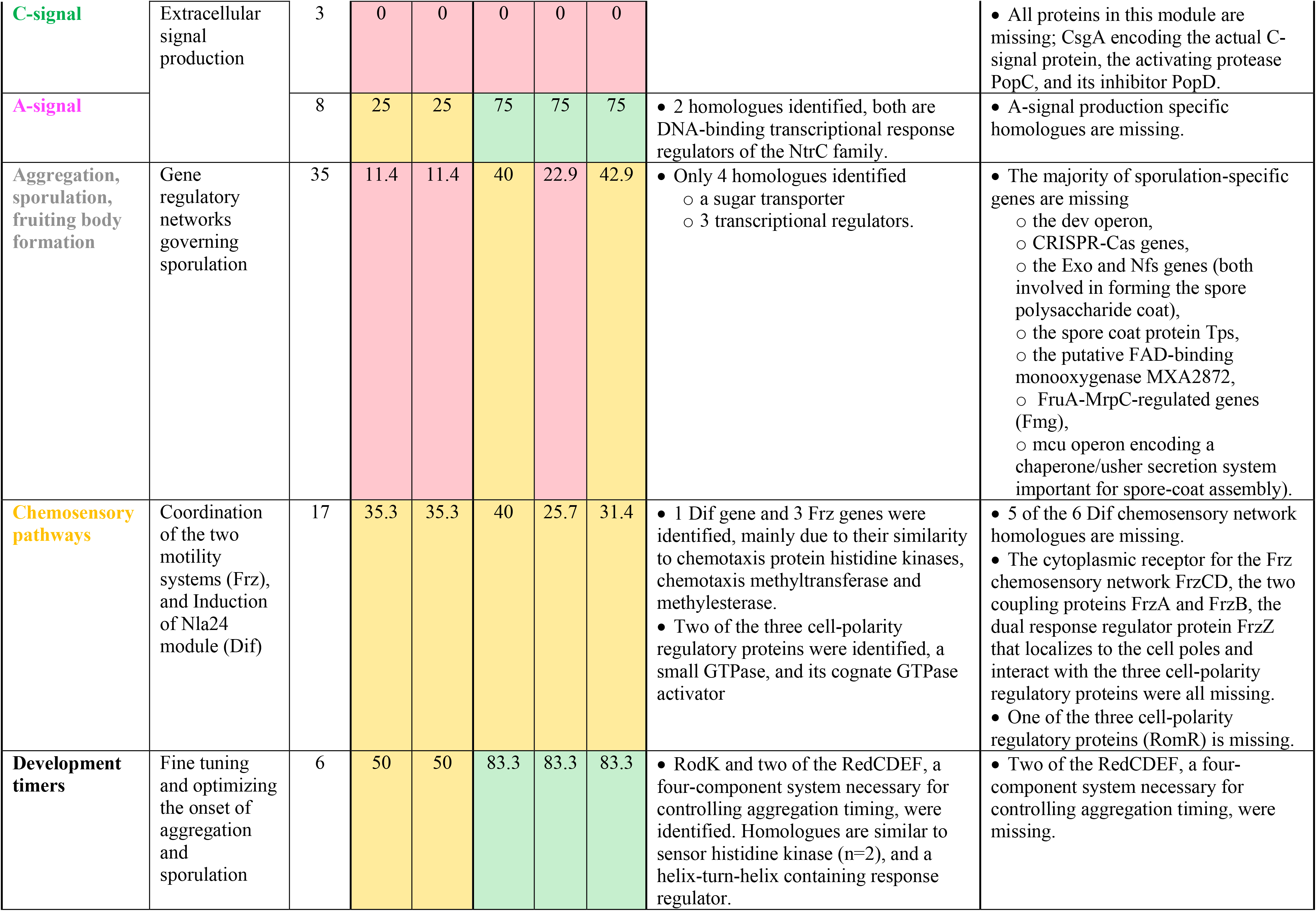
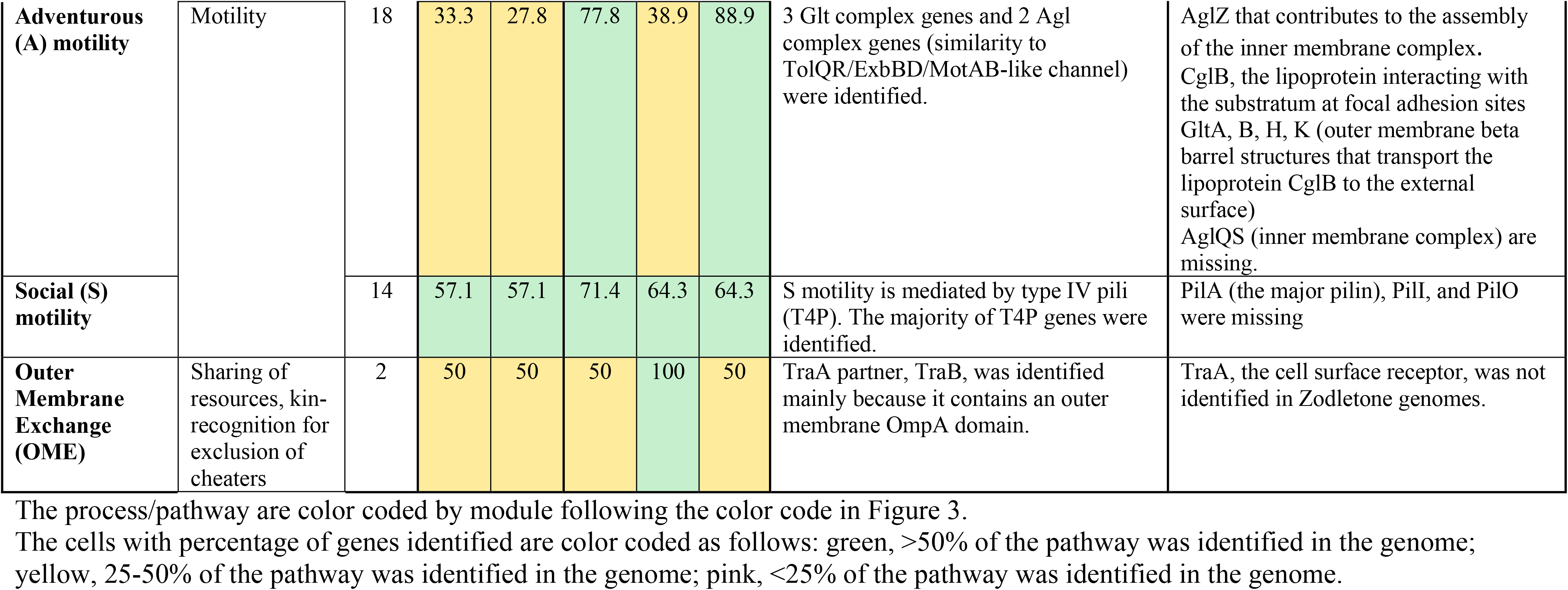
Comparative genome analysis of pathways implicated in social behavior between Zodletone and cultured Myxococcota.

### Machine learning approaches suggest absence of social behavior in Zodletone Myxococcota

It is important to note that most of the physiological, mutational, and transcriptomic studies on soil Myxococcota were conducted on the model organism *M. xanthus*, (Class Myxococcia), a relatively distant relative of Zodletone Myxococcota (Class Polyangia). Further, while genes for exopolysaccharide production, adventurous motility, extracellular signal production, and sporulation are highly specific, a large proportion of the gene regulatory network and developmental timing proteins governing aggregation, sporulation and fruiting body formation are homologues to signal transduction proteins involved in various cellular processes in a wide swaths of lineages, and are hence universally distributed within the bacterial world. Similarly, chemosensory network proteins in Myxococcota are homologues to a wide range of chemotaxis proteins. Indeed, in the genomes of the non-social *Anaeromyxobacter dehalogenes, Labilithrix luteola,* and *Vulgatibacter incomptus*, homologues for the highly specific extracellular signal production, and sporulation genes were not identified, but homologues for chemosensory networks and gene regulatory networks were found, attesting to the above caveats with the approach.

Therefore, as a complementary approach, we employed a machine learning technique to predict social behavior potential in Zodletone Myxococcota. The approach depends on first identifying, from the initial set of all genes in the genomes, a group of genes with assigned KO numbers in the genomes of known social Myxococcota that are absent from genomes of known non-social Myxococcota. These candidate genes were then used for model training using a Random Forest algorithm, and the constructed model was employed to predict the social behavior based on the genomic content of Zodletone Myxococcota genomes. The occurrence pattern of the list of 634 KOs (Dataset 1) selected for model training predicted non-social behavior for Zodletone Myxococcota lineages with a Matthew’s correlation coefficient of +1, confirming the patterns observed with the comparative genomics approach detailed above.

### Structural features and metabolic capacities

Structurally, Zodletone Myxococcota MAGs encoded determinants of Gram-negative cell walls (LPS biosynthesis and G-peptidoglycan structure), motility (flagellar assembly and type IV pili), pigmentation (carotenoid biosynthesis), chemotaxis, and rod-shape (MreBCD and RodA). The genomes also encoded Type III (partial) and type VI secretion systems. Such characteristics are similar to those displayed by vegetative cells of cultured Myxococcota (Figure 4a, Table S7).

**Figure 4.**
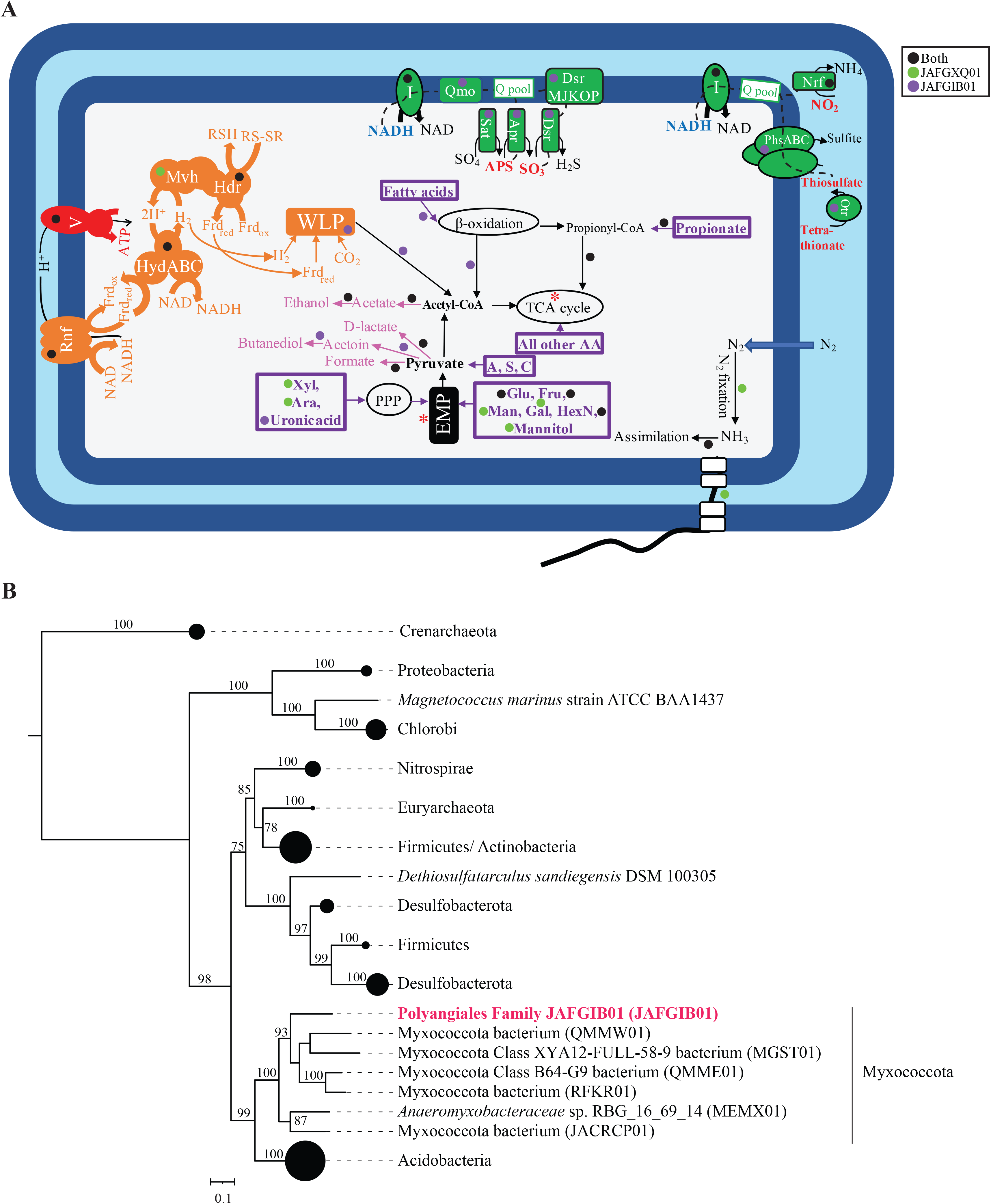
Metabolic capabilities predicted in Zodletone novel Myxococcota. (A) Cartoon depicting different metabolic capabilities encoded in the novel Zodletone genomes, with capabilities predicted for different lineages shown as colored circles (all orders, black; Order JAFGXQ01, green; Family JAFGIB01, purple). All substrates predicted to support growth are denoted in boldface purple text within thick purple boxes. Fermentation end products are shown in pink. Sites of substrate level phosphorylation are shown as red asterisks, as is the ATP synthase complex (V), while sites of proton extrusion to the periplasm are shown as black arrows. Electron transport components are shown in green. Paths of electron transfer are shown as dotted black lines that start at electron donors (shown in boldface blue text) ending at terminal electron acceptors (shown in boldface red text). Proton motive force generation, as well as electron carrier recycling pathways associated with the operation of the WLP are shown in orange. Abbreviations: Apr, the enzyme complex adenylylsulfate reductase [EC:1.8.99.2]; APS, adenylyl sulfate; Ara, arabinose; Dsr, dissimilatory sulfite reductase [EC:1.8.99.5]; EMP, Embden Meyerhoff Paranas pathway; Frd_ox/red_, Ferredoxin (oxidized/ reduced); Fru, fructose; Gal, galactose; Glu, glucose; Hdr, heterodisulfide reductase complex; HexN, hexosamines; HydABC, cytoplasmic [Fe Fe] hydrogenase; I, respiratory chain complex I; Man, mannose; Nrf, nitrite reductase (cytochrome c-552); Otr, octaheme reductase; PhsABC, thiosulfate reductase; PPP, pentose phosphate pathway; Pyr, pyruvate; Q pool, quinone pool; Qmo, quinone-interacting membrane-bound oxidoreductase complex; RNF, membrane-bound RNF complex; RSH/RS-SR, reduced/oxidized disulfide; TCA, tricarboxylic acid cycle; V, ATP synthase complex; WLP, Wood Ljungdahl pathway; Xyl, xylose. (B) Maximum likelihood phylogenetic tree based on the concatenated alignment of the alpha and beta subunits of dissimilatory sulfite reductase (DsrAB) from family JAFGIB01 (bold red text) in relation to reference sequences. Reference sequences from phyla other than Myxococcota are shown as wedged circles proportional to the number of sequences. Myxococcota references are shown with their GenBank assembly accession number in parentheses. Bootstrap support values are based on 100 replicates and are shown for nodes with >70% support.

Genomic analysis predicted key differences in anabolic capacities between Zodletone Myxococcota and cultured Myxococcota. Zodletone MAGs did not encode the capacity for glycogen or trehalose biosynthesis, both of which are biosynthesized and used as storage molecules by cultured Myxococcota, and shown to be essential for sporulation (71). Additionally, evidences for a glyoxylate shunt were missing from Zodletone MAGs. The glyoxylate shunt is employed by cultured Myxococcota to bypass CO_2_ loss and NADH production during the TCA cycle and drive the metabolism towards oxaloacetate in preparation for gluconeogenesis (72). Further, key differences in levels of amino acids auxotrophy were predicted, where Zodletone Myxococcota MAGs encoded capacities for biosynthesis of almost all amino acids, compared to the observed auxotrophy for branched chain amino acids (in 9 type species) and aromatic amino acids (in 5 type species) in cultured Myxococcota (73, 74). Such pattern reflects the dependence of cultured Myxococcota on proteins and amino acids as substrates (and hence their ready availability for biosynthetic purposes) as opposed to the lack of such capacity in Zodletone genomes, necessitating amino acid biosynthesis from metabolic precursors. Finally, while cultured Myxococcota are able to incorporate sulfur from sulfate as well as organic sources (e.g. taurine, alkane sulfonate, and dimethyl sulfone in 6 type species), and incorporate N from ammonia as well as organic sources (e.g. urea in 11 type species), such capacities for S or N incorporation from organic sources were not encoded in Zodletone Myxococcota MAGs (Table S7). Additionally, order JAFGXQ01 genomes encoded the capability to fix atmospheric nitrogen (Figure 4a, Table S7).

Genomic analysis also demonstrated multiple key differences in catabolic processes (substrate utilization patterns, respiratory capacities, electron recycling pathways, and ATP generation mechanisms) between Zodletone and model Myxococcota genomes. While most model Myxococcota (with the exception of *Sorangium cellulosum*) rely on amino acids and lipids as substrates and are poor carbohydrate consumers (71), Zodletone genomes encode a much lower number of amino acid degradation pathways (only 9, compared to 15 in type species), consistent with their observed limited proteolytic capabilities (Table S2). Instead, Zodletone Myxococcota appear to possess a more extensive carbohydrate degradation capacity (Tables 3, S7, Figure 4a), with pathways enabling the degradation of nine different sugars, sugar alcohols, sugar amines, and uronic acids encoded in their genomes. This is consistent with the possession of a wide range of polysaccharide-degrading CAZymes, as described above (Table S4). Finally, Zodletone order JAFGXQ01 genomes encoded an incomplete beta-oxidation pathway for long chain fatty acid degradation (Table 3, Figure 4a), a pathway commonly occurring in cultured Myxococcota to enable fatty acid consumption as the main carbon and energy source.

**Table 3.**
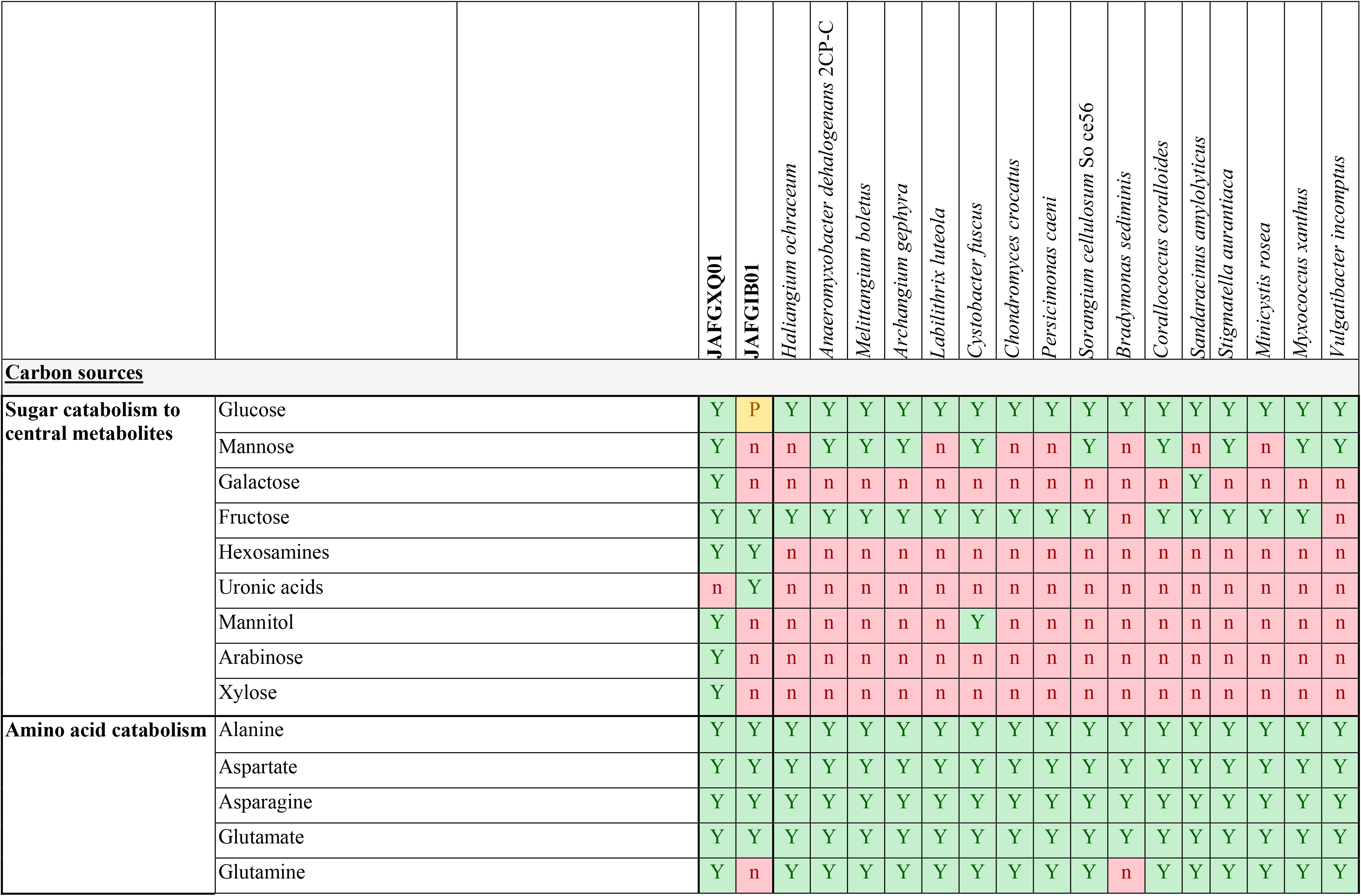
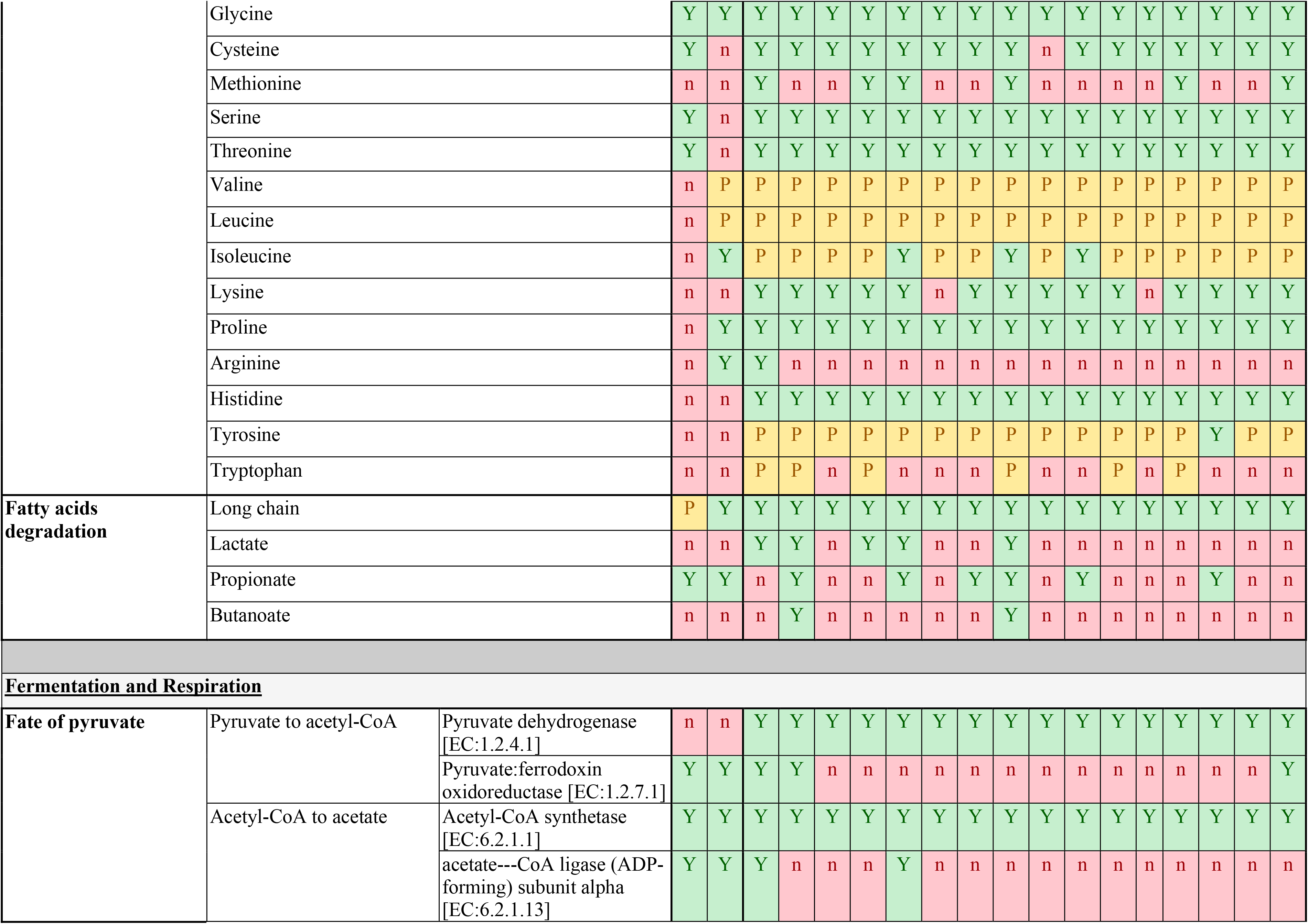
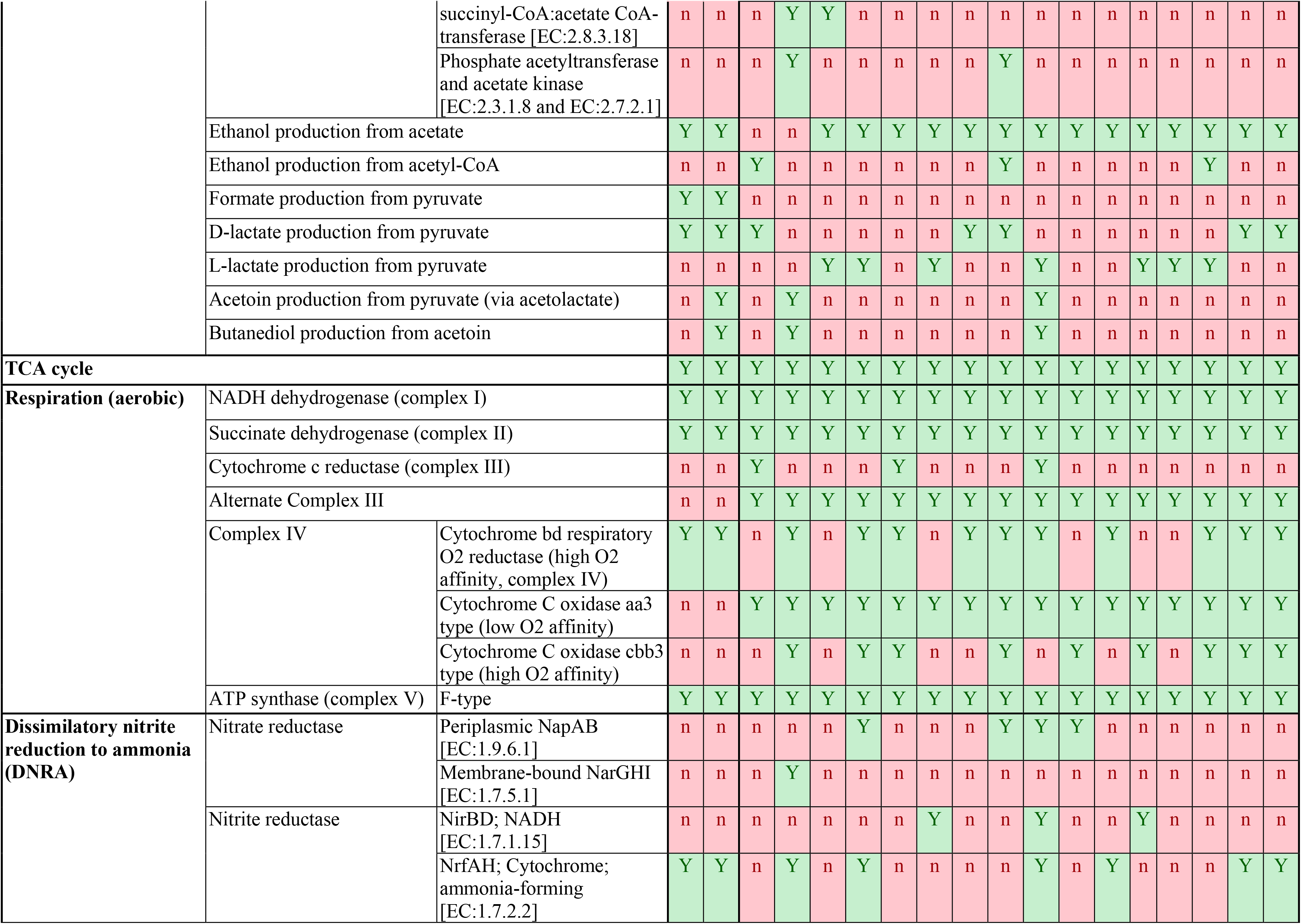
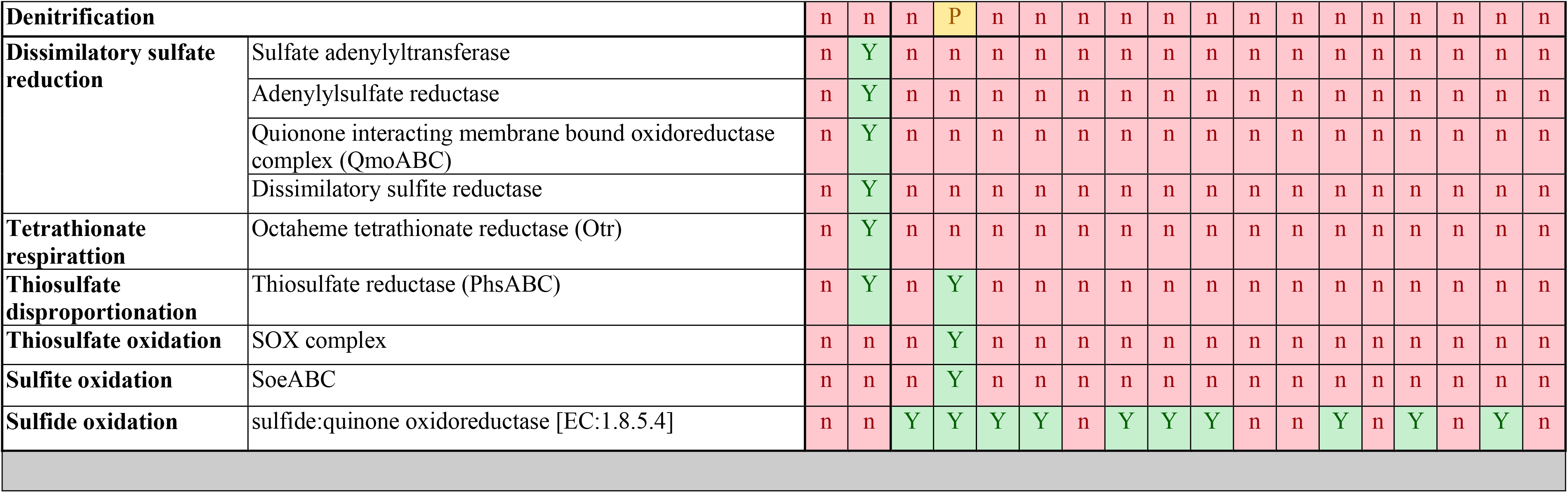
Catabolic capabilities deduced from genomic analysis of Zodletone Myxococcota in comparison to type species genomes available from KEGG organisms database. (Y, full pathway identified; n, full pathway missing; P, partial pathway identified).

All cultured Myxococcota (with the exception of the genus *Anaeromyxobacter*) are aerobic microorganisms. In contrast, Zodletone Myxococcota genomes lack key genes for aerobic respiration, specifically homologues for either complex III or alternative complex III, as well as the absence of homologues for the low affinity cytochrome oxidase aa3. High affinity cytochrome bd ubiquinol oxidase is encoded in Zodletone genomes, but could possibly be employed in detoxification of trace amounts of O_2_ present. Instead, the genomes encode genes enabling the utilization of nitrite as a terminal electron acceptor via the cytochrome-linked nitrite reductase NrfAH (Table 3, Figure 4a). Further, the single genome representative of novel family JAFGIB01 encodes a full dissimilatory sulfate reduction machinery including 3’-phosphoadenosine 5’-phosphosulfate synthase [Sat; EC:2.7.7.4 2.7.1.25] for sulfate activation to APS, adenylylsulfate reductase [AprAB; EC:1.8.99.2] for adenylyl sulfate reduction to sulfite, QmoABC for electron transfer, dissimilatory sulfite reductase [DsrAB; EC:1.8.99.5] and its co-substrate DsrC for dissimilatory sulfite reduction to sulfide, and the sulfite reduction-associated membrane complex DsrMKJOP for linking cytoplasmic sulfite reduction to energy conservation (Table 3, Figure 4a). JAFGIB01 genome also encoded octaheme tetrathionate reductase (*otr*) and thiosulfate reductase *phsABC,* suggesting the capability to utilize tetrathionate and thiosulfate as terminal electron acceptors in addition to sulfate (Figure 4a). This is the first genomic report of sulfur species respiration capability in the Myxococcota, and could possibly be a reflection of the sulfur and sulfide-rich Zodletone Spring environment from which the MAGs were binned. Phylogenetically, JAFGIB01 DsrAB were most closely affiliated to DsrAB sequences encountered in Acidobacteria genomes (Figure 4b) (34).

Besides respiration, additional pathways for electron disposal were identified in Zodletone Myxococcota genomes. These include fermentative processes for acetate, ethanol, and lactate production from pyruvate (Table 3, Figure 4a). As well, the genomes encoded a full Wood Ljungdahl pathway (WLP), most probably acting as an electron sink mechanism for re-oxidizing reduced ferredoxin, as previously noted in *Candidatus* Bipolaricaulota and Desulfobacterota genomes (35, 75). Finally, a possible additional mechanism for ATP production in Zodletone Myxococcota is the utilization of the RNF complex for re-oxidizing reduced ferredoxin at the expense of NAD, with the concomitant export of protons to the periplasm, generating a proton motive force that can drive ATP production via oxidative phosphorylation via the encoded F-type ATP synthase. Consistent with encoding RNF complex components, the genomes also encoded elements for electron carriers recycling including the cytoplasmic electron bifurcating mechanism HydABC. Analysis of Myxococcota type species genomes revealed absence of RNF complex components, HydABC electron bifurcation system, as well as the WLP pathway, consistent with a strictly aerobic mode of metabolism.

## Discussion

Our comparative genomics and machine learning analyses revealed severely curtailed machineries for predation and cellular differentiation in MAGs representing a novel order and a novel family of Myxococcota recovered from Zodletone spring (Figures 2, 3, Tables 2 and S6). Such results are in-agreement with a recent study that predicted absence of predation potential in MAGs/SAGs encompassing most of the publicly available, yet-uncultured Myxococcota (4). As such, a clear delineation exists between two phylogenetically and behaviorally distinct groups within the Myxococcota. The first encompasses aerobic top soil dwellers in classes Myxococcia and Polyangia that are characterized by possessing a highly sophisticated machinery enabling predation and cellular differentiation behaviors. Few freshwater and marine strains possessing such capacities have been reported, but their presence has been attributed to air and dust transport from neighboring soils (15, 16). Members of this group could readily be obtained in pure cultures. The second group encompasses phylogenetically distinct families and orders within the classes Myxococcia and Polyangia (including Zodletone MAGs), as well a few yet-uncultured Myxococcota classes. These lineages are almost invariably encountered within non-soil habitats (e.g. freshwater, marine, host-associated, and engineered ecosystems), and appear to lack the capacity for predation and social differentiation. Most of these lineages are currently uncultured, with the exception of members of the genus *Anaeromyxobacter* (76).

We argue that such patterns could provide important clues on the evolution of social behavior in the Myxococcota, when considered in light of our understanding of the history of soil formation and the rise of atmospheric oxygen in the atmosphere. Soil formation and transition from barren crusts to current soil orders through organic matter deposition and transformation has been enabled by the evolution of lichen associations, plant terrestrialization, formation of mycorrhizal association, and subsequent colonization by soil microfauna. All such processes are mediated by aerobic organisms (algae, fungi, plants, fauna), and hence was possible only after the accumulation of oxygen to levels comparable to current values in the atmosphere (approximately 500-600 Mya (77)). The formation of soil structures as a new and organic-rich habitat has certainly spurred multiple evolutionary processes for enabling terrestrial adaptation within the microbial world. Various processes have been reported in multiple soil-prevalent lineages, from CAZymes and BGCs acquisition in the Acidobacteria (34, 78), to acquisition of stress tolerance, adherence, and regulatory genes in the ammonia-oxidizing archaea (79). Here, it appears that the development of predation and cellular differentiation machineries has enabled the Myxococcota to assume an apex predator niche and imparted them with strong survival capacities in soil, respectively. Indeed, as previously noted, the ecological success of social Myxococcota in soils appears to be in stark contrast to their rarity and low relative abundance of non-social Myxococcota in other habitats (80).

Evolution of beneficial trait(s) in a single lineage of Myxococcota in soil could be propagated to the broader soil Myxococcota community through intra-clade, habitat-specific horizontal gene transfer (HGT), resulting in the observed checkered distribution pattern, where social behavior is observed only in specific families within the classes Myxococcia and Polyangia. HGT between closely related taxa is a well-established phenomenon (81) that has been widely documented, e.g. in mediating the spread of antibiotic resistance in related clinical strains (82, 83). The barrier for HGT within closely related taxa is predictably lower, given expected similarity in codon usage pattern, GC content, restriction enzyme machinery, and overall genome architecture between donor and recipient strains. Similarly, physical proximity in the same habitat is seen as a facilitator of genetic exchange through HGT (84–86).

In conclusion, our results strongly indicate that anaerobic Myxococcota do not posses the capacity for typical social behavior, and display distinct structural, anabolic, and catabolic differences when compared to model aerobic Myxococcota. We document their dependence on fermentation and/or nitrite and sulfate-reduction for energy generation, as well as their preference for polysaccharide metabolism over protein, amino acids, and lipid metabolism. We further propose that such differences strongly underscore the importance of niche differentiation in shaping the evolutionary trajectory of the Myxococcota, and suggest soil formation as a strong driver for developing social behavior in this lineage.

## Acknowledgments

We thank Dr. David W. Waite at the Australian Centre for Ecogenomics, School of Chemistry and Molecular Biosciences, University of Queensland, for helpful discussions regarding utilization of machine learning approaches for trait prediction. This work was supported by NSF grant 2016423 to NHY and MSE.

